## Supplementary Figures and Tables for "CemR atypical response regulator impacts energy conversion in *Campylobacteria*"

### Fig. S1

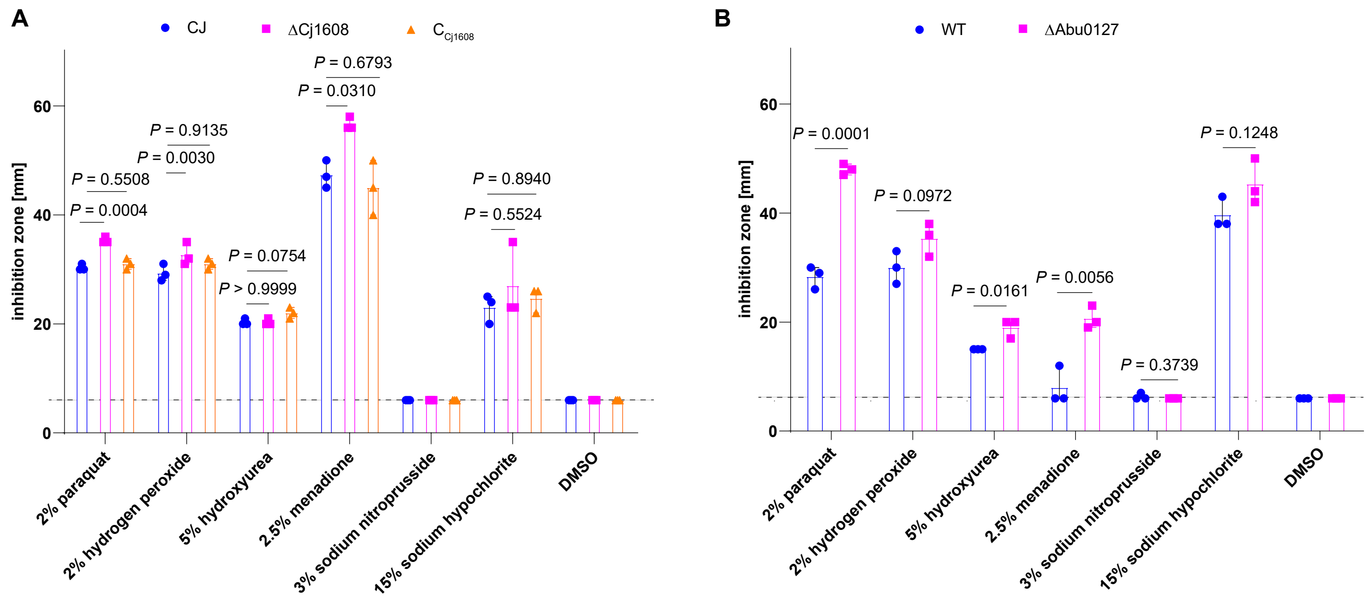
A
CJ
ΔCj1608
C Cj1608
inhibition zone [mm]
P = 0.5508
P = 0.0004
P = 0.9135
P = 0.0030
P = 0.0754
P > 0.9999
P = 0.6793
P = 0.0310
P = 0.8940
P = 0.5524
2% paraquat
2% hydrogen peroxide
5% hydroxyurea
2.5% menadione
3% sodium nitroprusside
15% sodium hypochlorite
DMSO
B
WT
ΔAbu0127
P = 0.0001
P = 0.0972
P = 0.0161
P = 0.0056
P = 0.3739
P = 0.1248

### Fig. S2

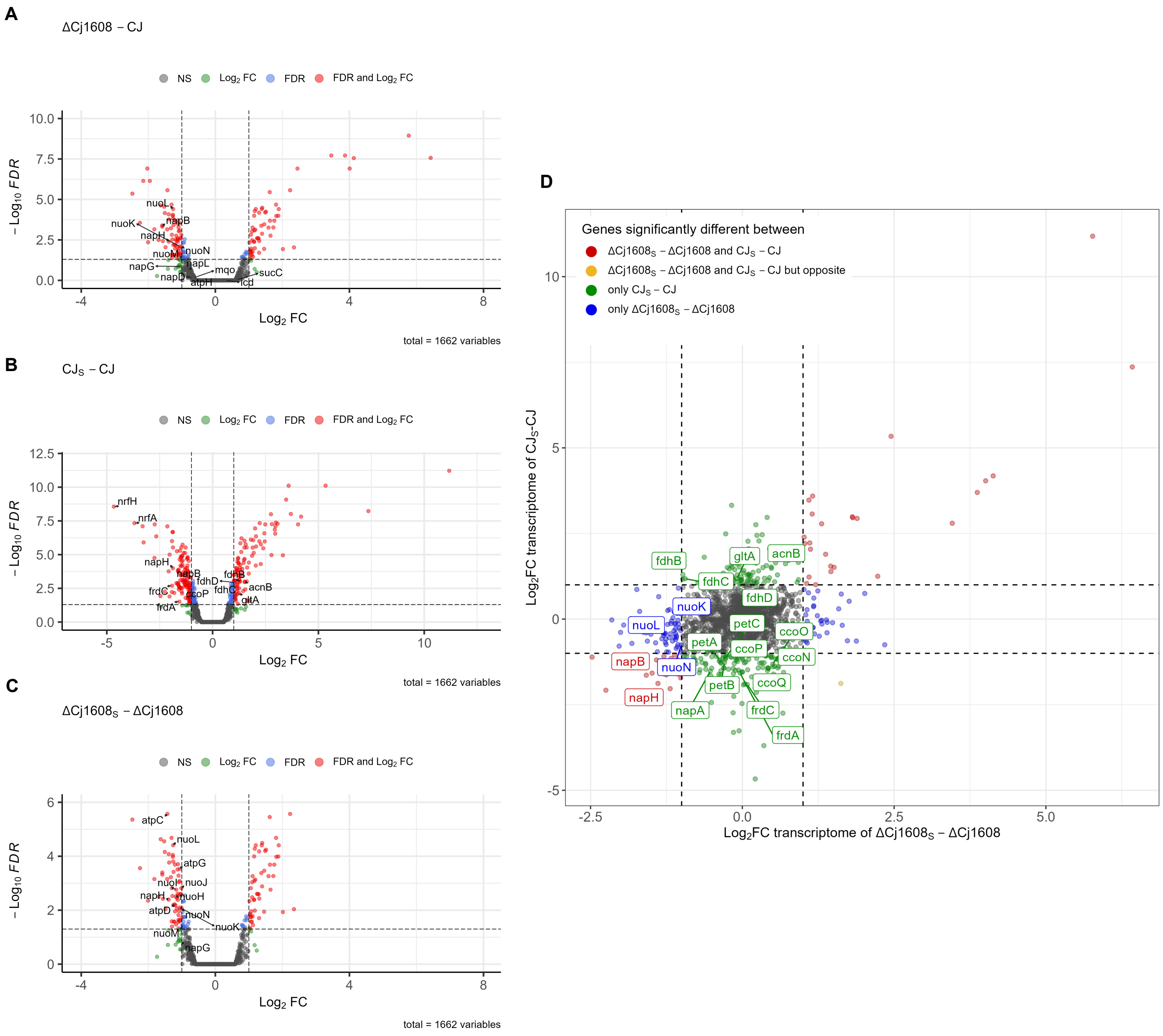
A
ΔCj1608 − CJ
NS
Log2 FC
FDR
FDR and Log2 FC
− Log10 FDR
Log2 FC
total = 1662 variables
nuoL
napB
nuoK
napH
nuoN
nuoM
napL
napG
mqo
napD
atpH
sucC
icd
B
CJS − CJ
nrfH
nrfA
napH
napB
fdhD
fdhB
frdC
fdhC
acnB
ccoP
gltA
frdA
C
ΔCj1608S − ΔCj1608
atpC
nuoL
atpG
nuoI
nuoJ
napH
nuoH
atpD
nuoN
nuoK
nuoM
napG
D
Genes significantly different between
ΔCj1608S − ΔCj1608 and CJS − CJ
ΔCj1608S − ΔCj1608 and CJS − CJ but opposite
only CJS − CJ
only ΔCj1608S − ΔCj1608
Log2FC transcriptome of CJS-CJ
Log2FC transcriptome of ΔCj1608S − ΔCj1608
fdhB
gltA
acnB
fdhC
nuoK
fdhD
nuoL
petC
ccoO
petA
ccoP
ccoN
napB
nuoN
petB
ccoQ
napH
napA
frdC
frdA

### Fig. S3

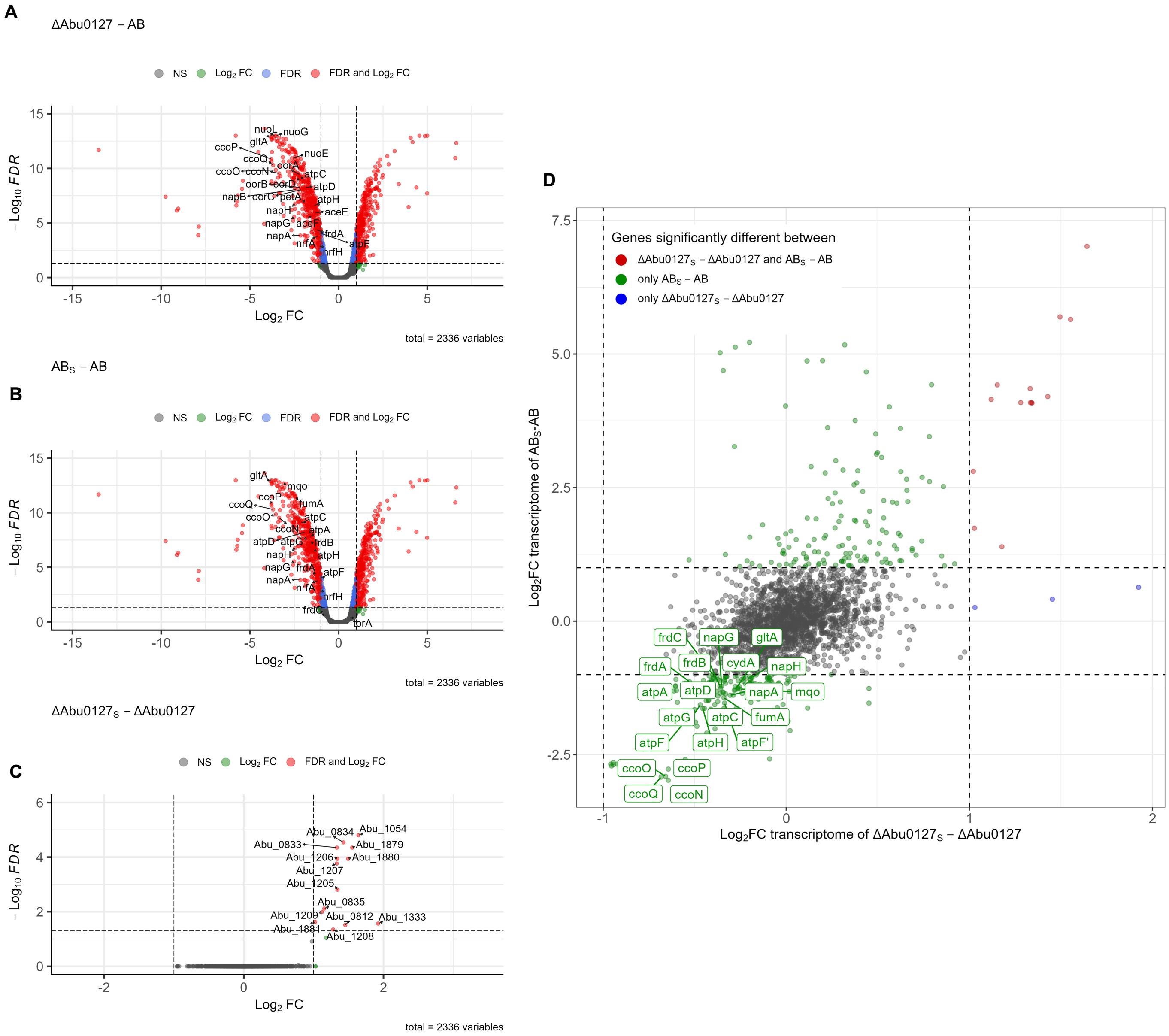

A
ΔAbu0127 − AB
NS
Log2 FC
FDR
FDR and Log2 FC
−Log10 FDR
Log2 FC
total = 2336 variables
nuoL
nuoG
gltA
ccoP
ccoQ
nuoE
oorA
ccoO
ccoN
atpC
oorB
oorD
atpD
napB
oorC
petA
atpH
napH
aceE
napG
aceF
napA
frdA
nrfA
atpF
nrfH
B
ABS − AB
NS
Log2 FC
FDR
FDR and Log2 FC
−Log10 FDR
Log2 FC
total = 2336 variables
gltA
mqo
ccoP
ccoQ
fumA
ccoO
atpC
ccoN
atpA
atpD
atpG
frdB
napH
atpH
napG
frdA
atpF
napA
nrfA
nrfH
frdC
torA
C
ΔAbu0127S − ΔAbu0127
NS
Log2 FC
FDR and Log2 FC
−Log10 FDR
Log2 FC
total = 2336 variables
Abu_0834
Abu_1054
Abu_0833
Abu_1879
Abu_1206
Abu_1880
Abu_1207
Abu_1205
Abu_0835
Abu_1209
Abu_0812
Abu_1333
Abu_1881
Abu_1208
D
Genes significantly different between
ΔAbu0127S − ΔAbu0127 and ABS − AB
only ABS − AB
only ΔAbu0127S − ΔAbu0127
Log2FC transcriptome of ABS-AB
Log2FC transcriptome of ΔAbu0127S − ΔAbu0127
frdC
napG
gltA
frdA
frdB
cydA
napH
atpA
atpD
napA
mqo
atpG
atpC
fumA
atpF
atpH
atpF'
ccoO
ccoP
ccoQ
ccoN

**Fig. S4**

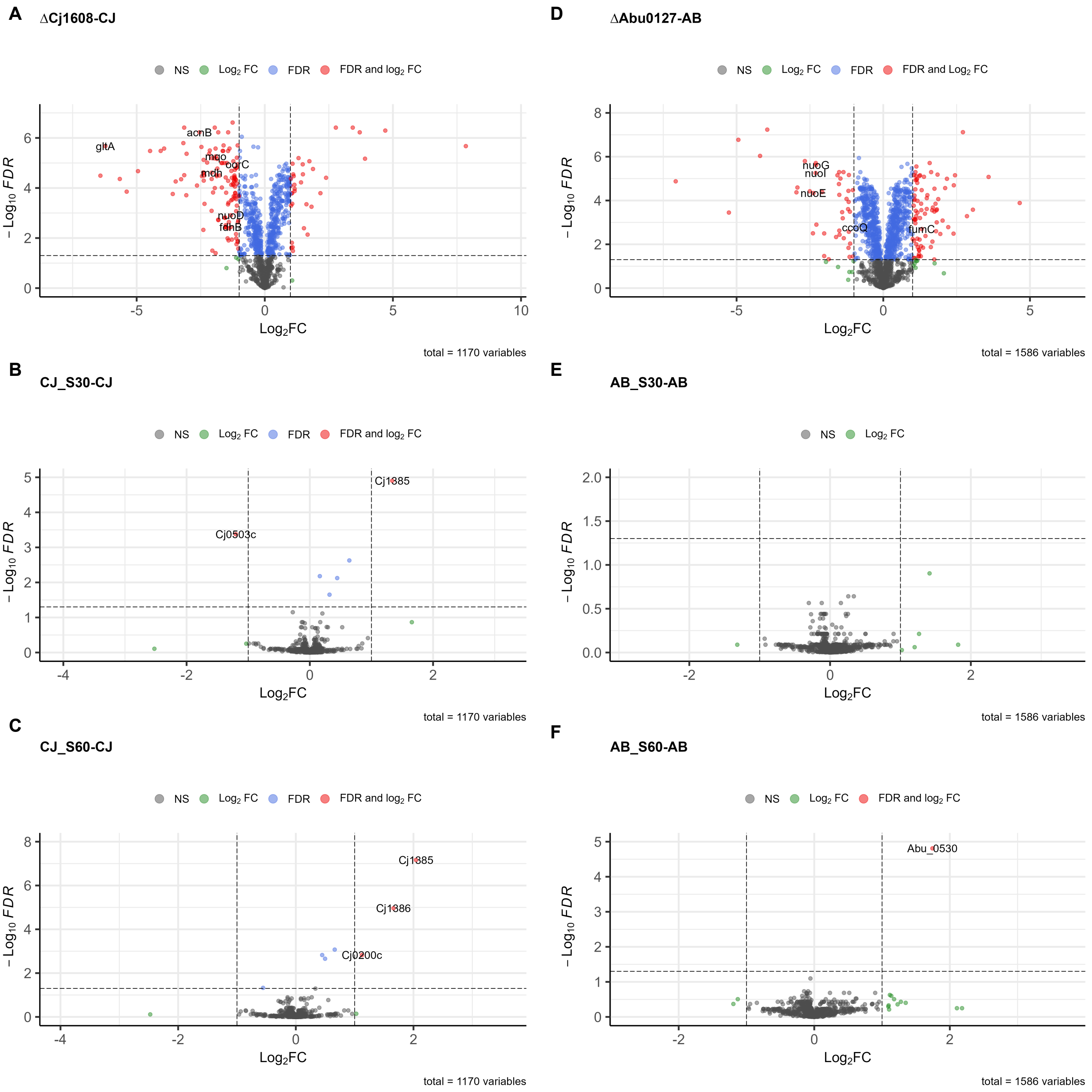
A
ΔCj1608-CJ
NS Log2 FC FDR FDR and log2 FC
gltA
acnB
mqo
oorC
mdh
nuoD
fdhB
− Log10 FDR
Log2FC
total = 1170 variables
B
CJ_S30-CJ
NS Log2 FC FDR FDR and log2 FC
Cj1885
Cj0503c
− Log10 FDR
Log2FC
total = 1170 variables
C
CJ_S60-CJ
NS Log2 FC FDR FDR and log2 FC
Cj1885
Cj1886
Cj0200c
− Log10 FDR
Log2FC
total = 1170 variables
D
ΔAbu0127-AB
NS Log2 FC FDR FDR and Log2 FC
nuoG
nuoI
nuoE
ccoQ
fumC
− Log10 FDR
Log2FC
total = 1586 variables
E
AB_S30-AB
NS Log2 FC
− Log10 FDR
Log2FC
total = 1586 variables
F
AB_S60-AB
NS Log2 FC FDR and Log2 FC
Abu_0530
− Log10 FDR
Log2FC
total = 1586 variables

**Fig. S5**

**A**

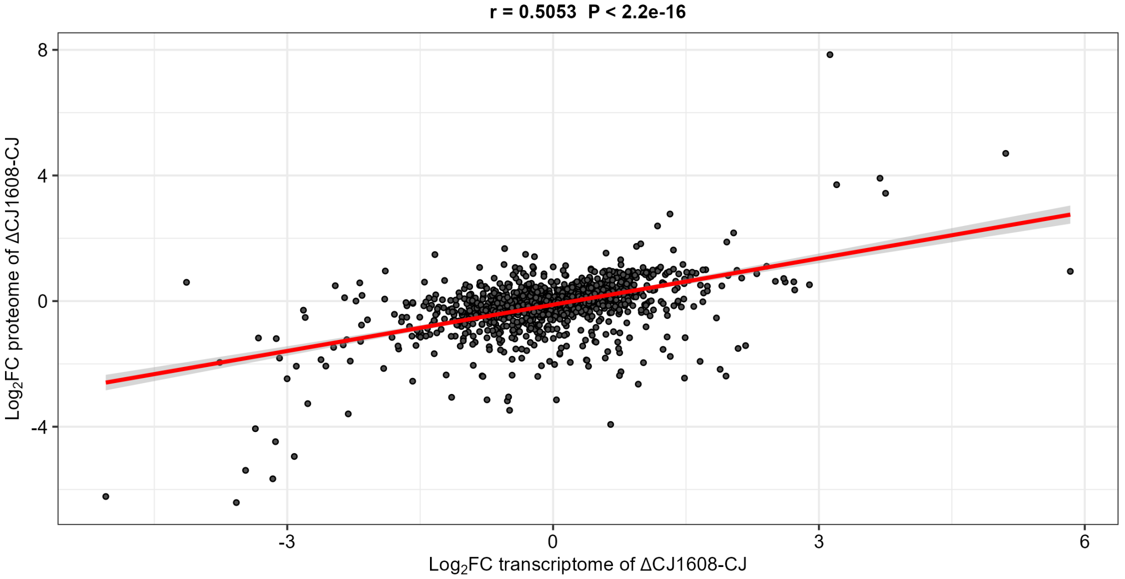

r = 0.5053 P < 2.2e-16
8
4
0
-4
-3
0
3
6
Log2FC proteome of ΔCJ1608-CJ
Log2FC transcriptome of ΔCJ1608-CJ


**B**

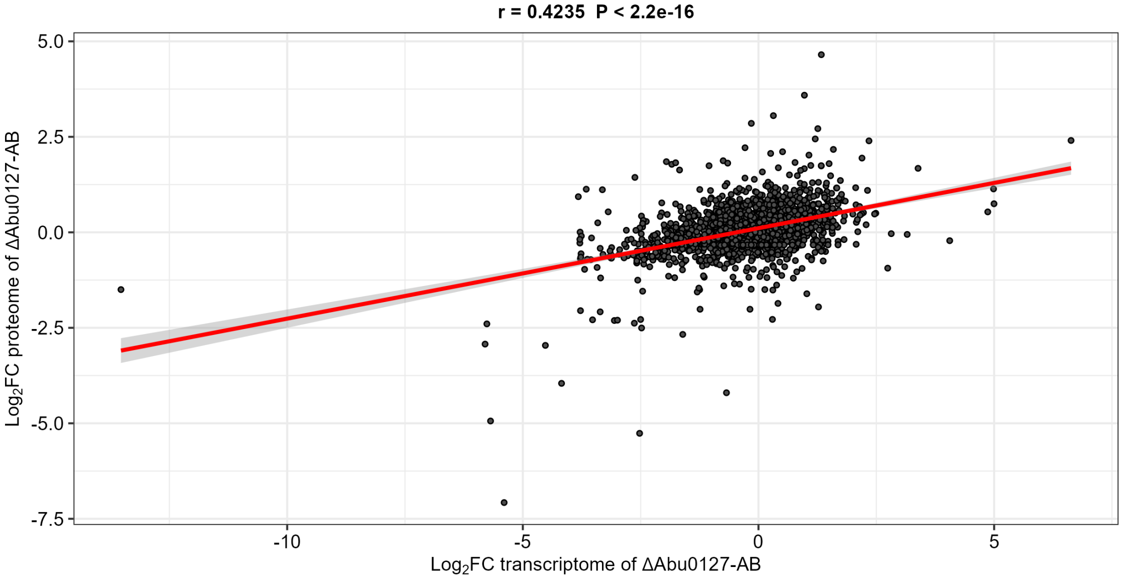

r = 0.4235 P < 2.2e-16
5.0
2.5
0.0
-2.5
-5.0
-7.5
-10
-5
0
5
Log2FC proteome of ΔAbu0127-AB
Log2FC transcriptome of ΔAbu0127-AB

### Fig. S6

**A**

RNA-seq

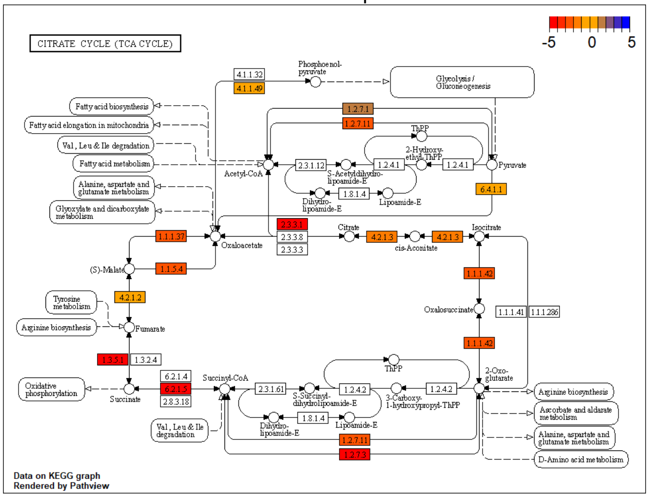


**B**

LC-MS/MS

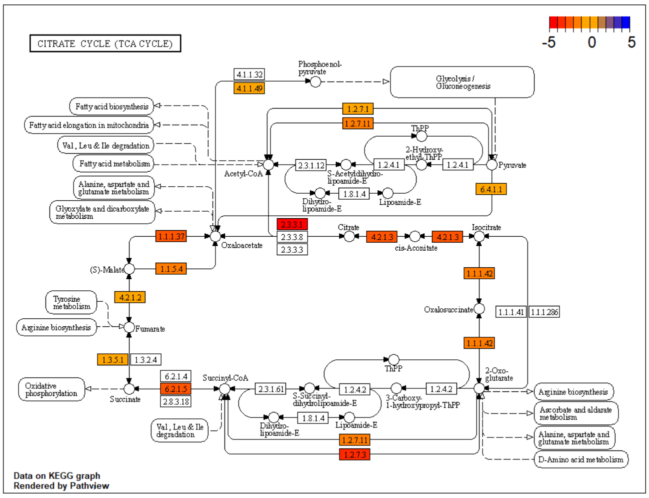

### Fig. S7

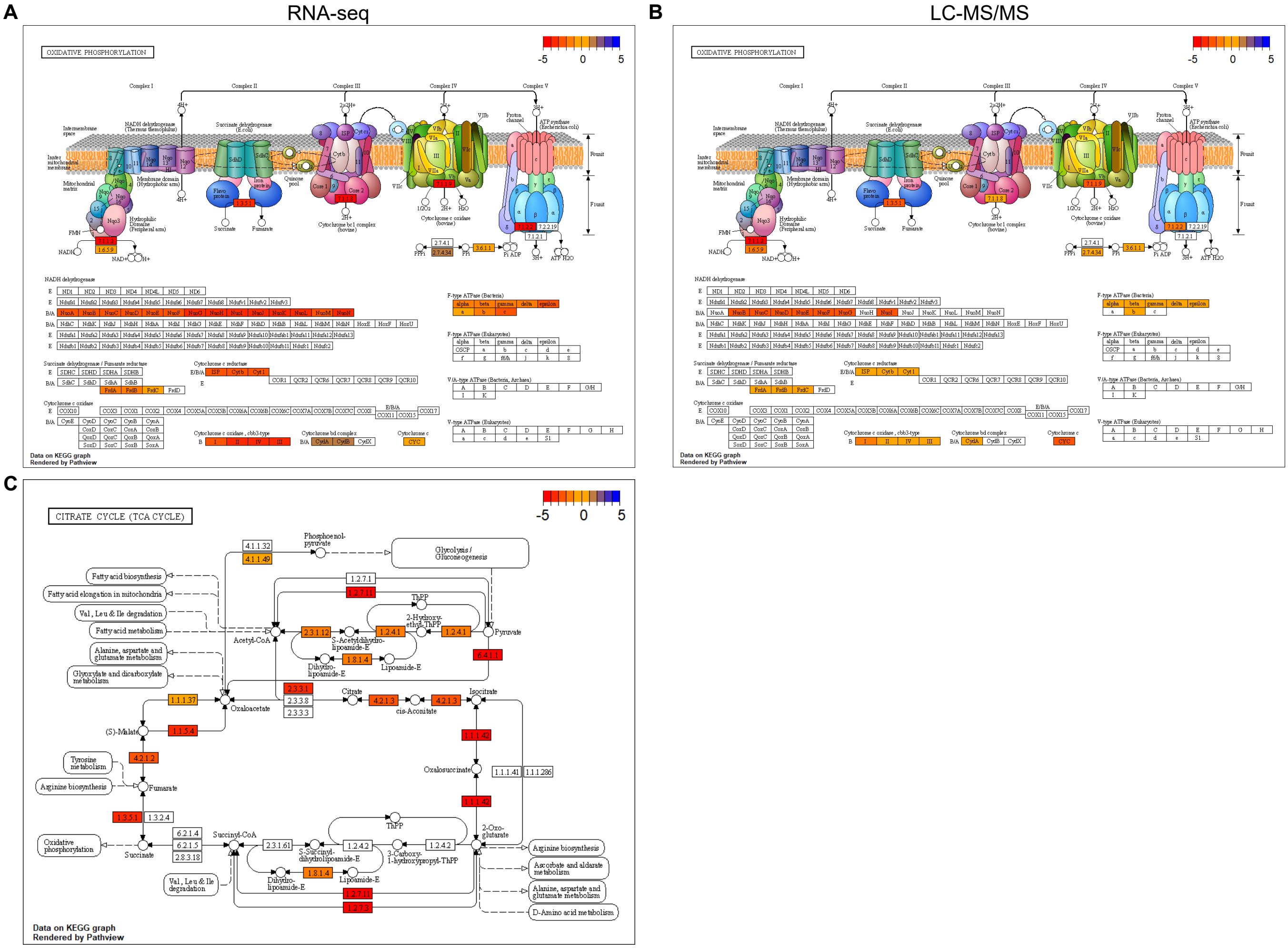
A
RNA-seq
B
LC-MS/MS
C
OXIDATIVE PHOSPHORYLATION
-5
0
5
Data on KEGG graph
Rendered by Pathview
CITRATE CYCLE (TCA CYCLE)
Phosphoenol-pyruvate
Glycolysis / Gluconeogenesis
Fatty acid biosynthesis
Fatty acid elongation in mitochondria
Val, Leu & Ile degradation
Fatty acid metabolism
Alanine, aspartate and glutamate metabolism
Glyoxylate and dicarboxylate metabolism
Acetyl-CoA
Pyruvate
Oxaloacetate
Citrate
cis-Aconitate
Isocitrate
(S)-Malate
Tyrosine metabolism
Arginine biosynthesis
Fumarate
Oxalosuccinate
Succinate
Succinyl-CoA
Oxidative phosphorylation
2-Oxo-glutarate
Ascorbate and aldarate metabolism
D-Amino acid metabolism

#### Fig. S8

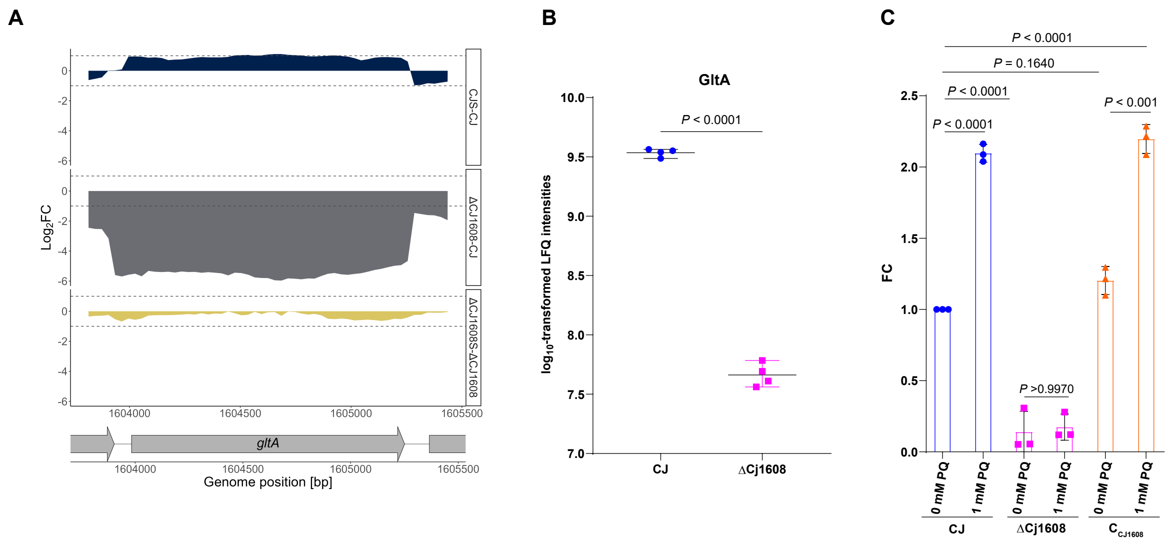
A
Log2FC
CJS-CJ
ΔCJ1608-CJ
ΔCJ1608S-ΔCJ1608
gltA
Genome position [bp]
B
GltA
P < 0.0001
log10-transformed LFQ intensities
CJ
ΔCj1608
C
P < 0.0001
P = 0.1640
P < 0.0001
P < 0.0001
P < 0.001
P >0.9970
FC
0 mM PQ
1 mM PQ
CJ
ΔCj1608
CCJ1608

### Fig. S9

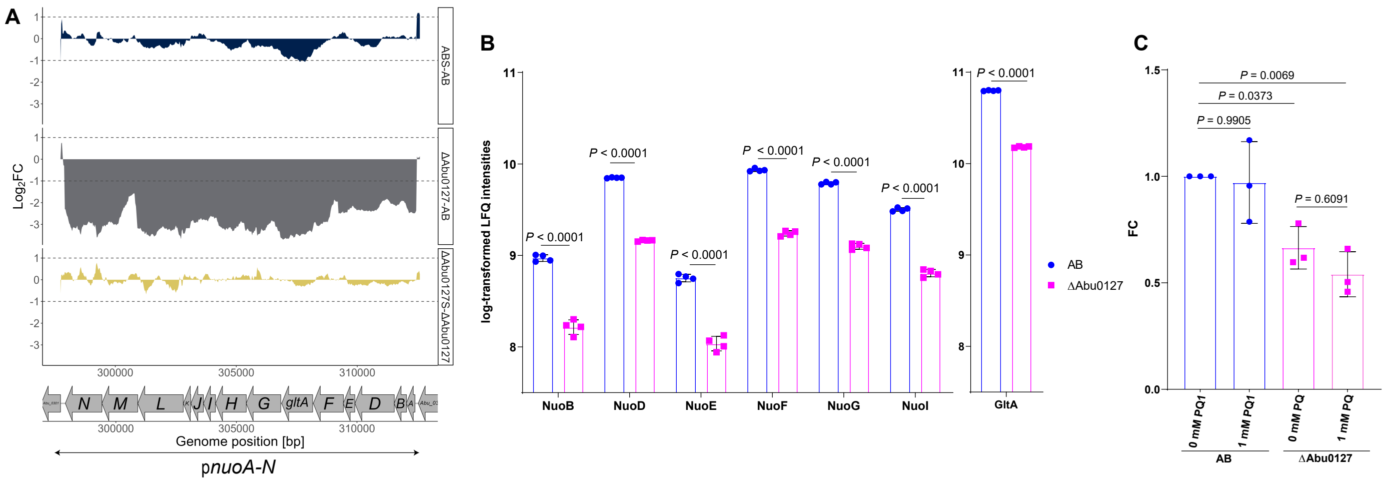

A
ABS-AB
ΔAbu0127-AB
ΔAbu0127S-ΔAbu0127
Log2FC
300000
305000
310000
N
M
L
J
I
H
G
gltA
F
E
D
B
A
Genome position [bp]
pnuoA-N
B
log-transformed LFQ intensities
P < 0.0001
NuoB
NuoD
NuoE
NuoF
NuoG
NuoI
GltA
AB
ΔAbu0127
C
FC
P = 0.0069
P = 0.0373
P = 0.9905
P = 0.6091
0 mM PQ1
1 mM PQ1
0 mM PQ
1 mM PQ
AB
ΔAbu0127

#### Fig. S10

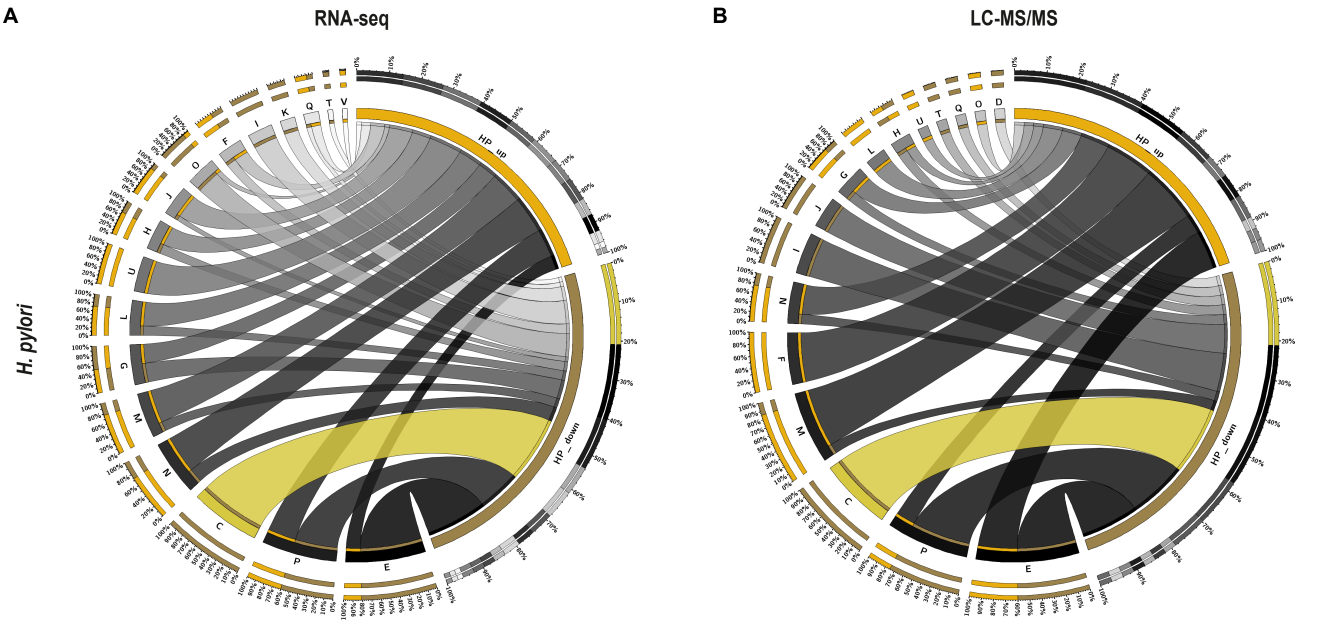


**C** - energy production and conversion, **D** - cell cycle control, cell division, chromosome partitioning, **E** - amino acid transport and metabolism, **F** - nucleotide transport and metabolism, **G** - carbohydrate transport and metabolism, **H** - coenzyme transport and metabolism, **I** - lipid transport and metabolism, **J** - translation, ribosomal structure and biogenesis, **K** - transcription, **L** - replication, recombination and repair, **M** - cell wall/membrane/envelope biogenesis, **N** - cell motility, **O** - posttranslational modification, protein turnover, chaperones , **P** - inorganic ion transport and metabolism, **Q** - secondary metabolites biosynthesis, transport and catabolism, **T** - signal transduction mechanisms, **U** - intracellular trafficking, secretion, and vesicular transport, **V** - defense mechanisms

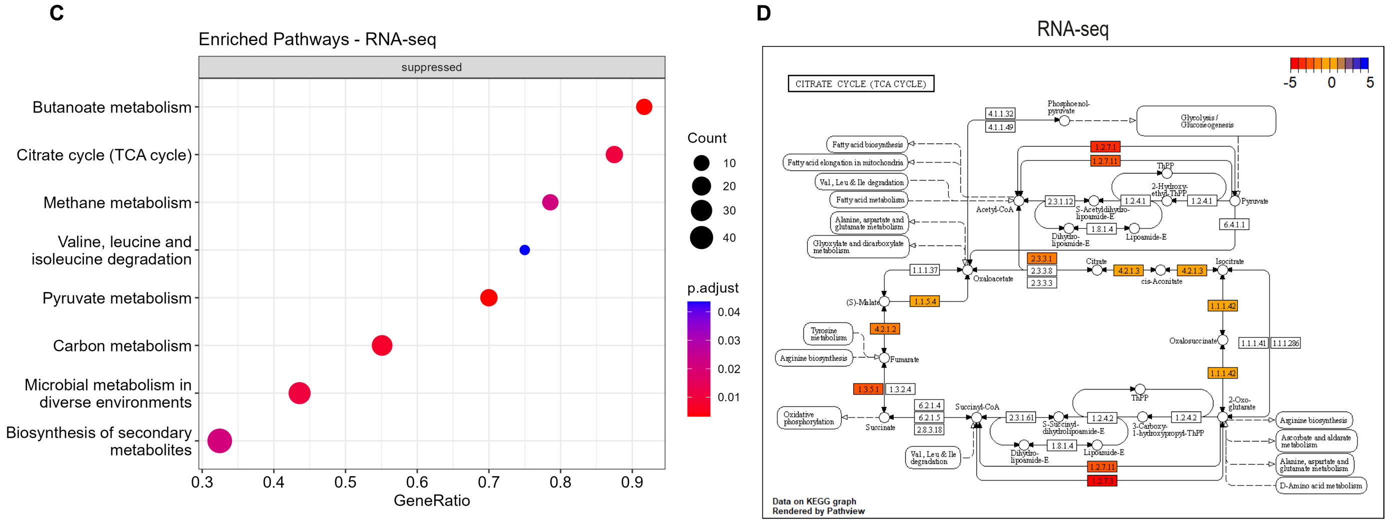

#### Fig. S11

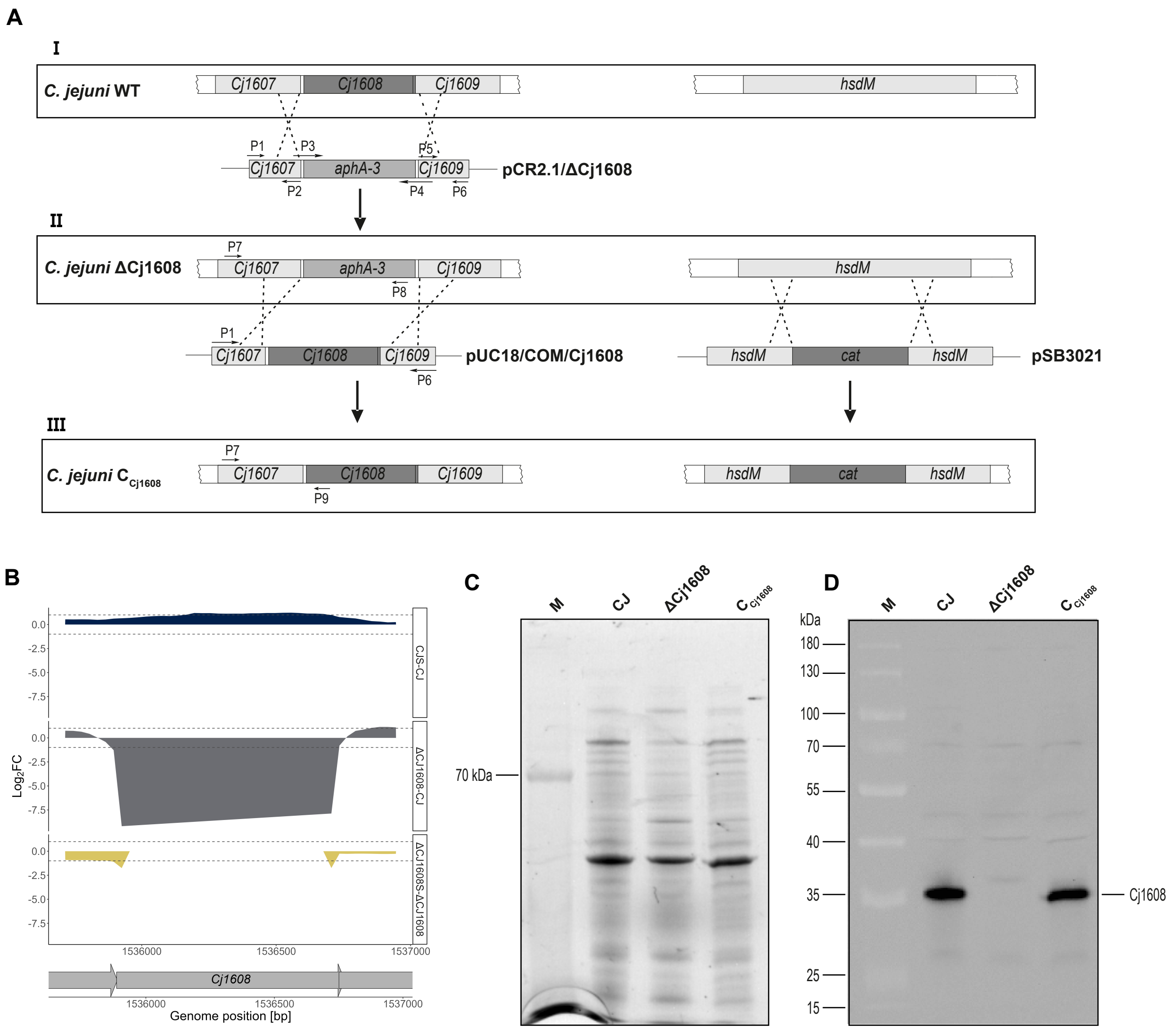

A
I
C. jejuni WT
Cj1607
Cj1608
Cj1609
hsdM
P1
P3
P5
P2
P4
P6
Cj1607
aphA-3
Cj1609
pCR2.1/ΔCj1608
II
C. jejuni ΔCj1608
P7
Cj1607
aphA-3
Cj1609
P8
hsdM
P1
Cj1607
Cj1608
Cj1609
P6
pUC18/COM/Cj1608
hsdM
cat
hsdM
pSB3021
III
C. jejuni C Cj1608
P7
Cj1607
Cj1608
Cj1609
P9
hsdM
cat
hsdM
B
Log2FC
0.0
-2.5
-5.0
-7.5
CJS-CJ
ΔCJ1608-CJ
ΔCJ1608S-ΔCJ1608
1536000
1536500
1537000
Cj1608
Genome position [bp]
C
M
CJ
ΔCj1608
C Cj1608
70 kDa
D
M
CJ
ΔCj1608
C Cj1608
kDa
180
130
100
70
55
40
35
25
15
Cj1608

### Fig. S12

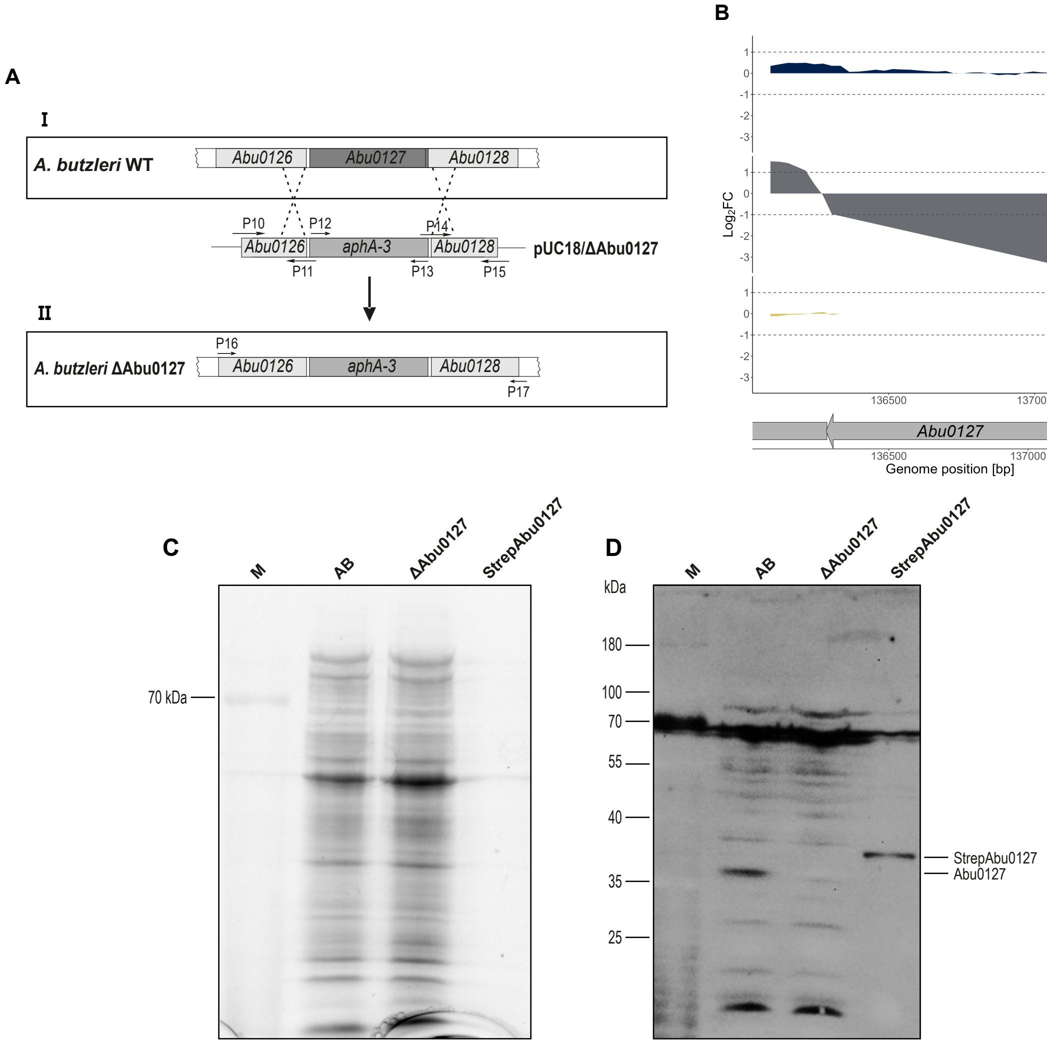
A
I
A. butzleri WT
Abu0126
Abu0127
Abu0128
P10
P12
P14
Abu0126
aphA-3
Abu0128
pUC18/ΔAbu0127
P11
P13
P15
II
P16
A. butzleri ΔAbu0127
Abu0126
aphA-3
Abu0128
P17
B
Log2FC
ABS-AB
ΔAbu0127-AB
ΔAbu0127S-ΔAbu0127
136500
137000
Abu0127
Genome position [bp]
C
M
AB
ΔAbu0127
StrepAbu0127
70 kDa
D
kDa
180
100
70
55
40
35
25
StrepAbu0127
Abu0127

**Fig. S13**

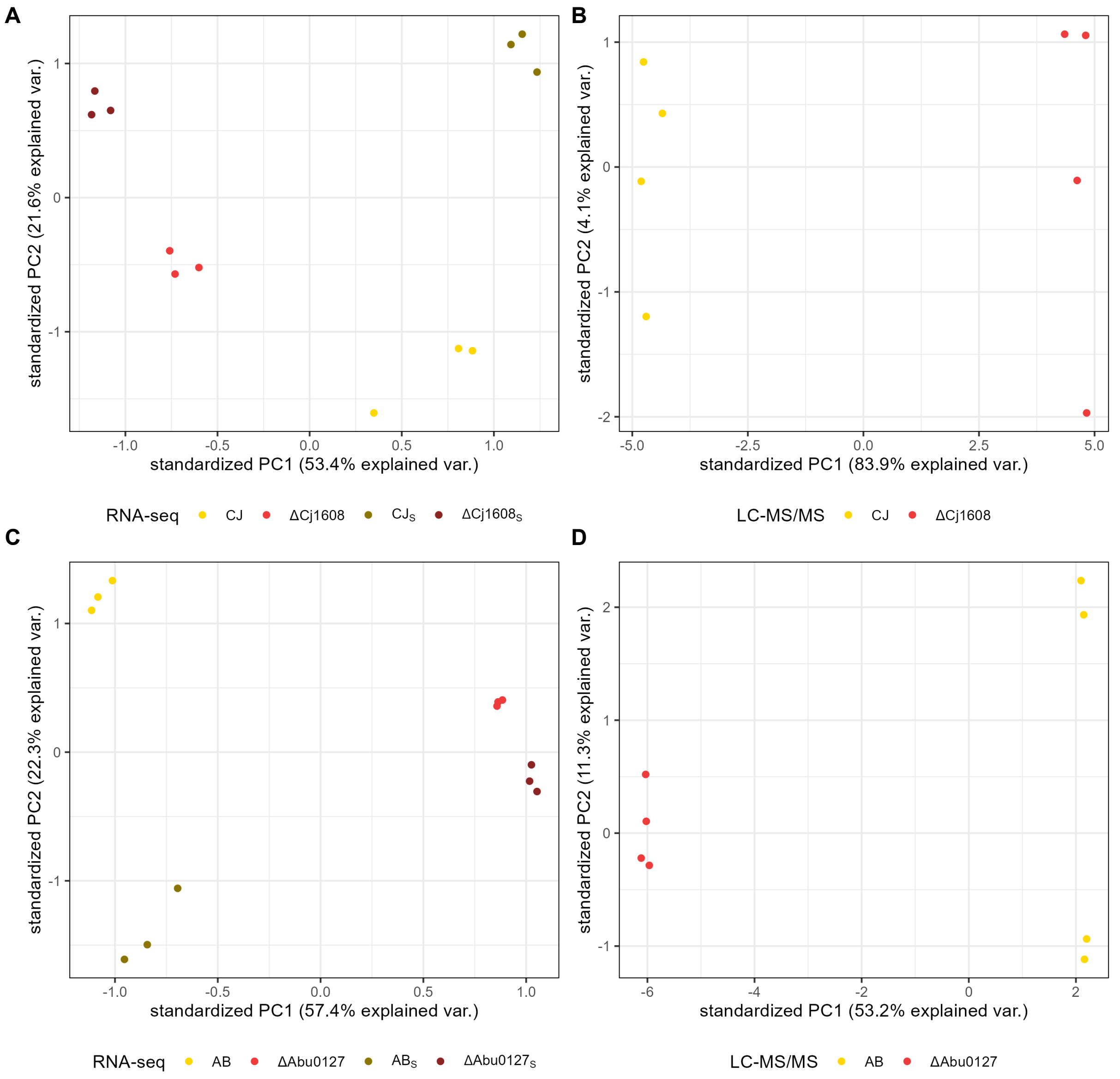
A
B
C
D
standardized PC1 (53.4% explained var.)
standardized PC2 (21.6% explained var.)
standardized PC1 (83.9% explained var.)
standardized PC2 (4.1% explained var.)
RNA-seq CJ ΔCj1608 CJ$_S$ ΔCj1608$_S$
LC-MS/MS CJ ΔCj1608
standardized PC1 (57.4% explained var.)
standardized PC2 (22.3% explained var.)
standardized PC1 (53.2% explained var.)
standardized PC2 (11.3% explained var.)
RNA-seq AB ΔAbu0127 AB$_S$ ΔAbu0127$_S$
LC-MS/MS AB ΔAbu0127

**Table S1: Strains, plasmids and proteins used in this study.**

| Strain | Relevant features | Reference/source |
|---|---|---|
| *E. coli* DH5α | *supE44*, *hsdR17*, *recA1*, *endA1*, *gyrA1*, *gyrA96*, *thi-1*, *relA1* | [1] |
| *E. coli* BL21 | F-, *ompT*, *hsdS (rB-*, *mB-)*, *gal*, *dcm* | GE Healthcare |
| *C. jejuni* NCTC 11168 | Parental strain | [2] |
| *C. jejuni* NCTC 11168 ΔCj1608 | Δ*Cj1608*::*aphA-3*; 11168 with *Cj1608* exchanged to *aphA-3* cassette | This study |
| *C. jejuni* NCTC 11168 $C_{Cj1608}$ | (Δ*Cj1608*::*aphA-3*)::*Cj1608*, *hdsM*::*cat*; 11168 ΔCj1608 in which *aphA-3* was exchanged to *Cj1608* and *cat* cassette was inserted in to *hsdM* gene | This study |
| *A. butzleri* RM4018 | Parental strain | [3], DSMZ-German Collection of Microorganisms and Cell Cultures GmbH |
| *A. butzleri* RM4018 ΔAbu0127 | Δ*Abu0127*::*aphA-3*; Abu0127 exchanged to *aphA-3* cassette | This study |

| Plasmid | Relevant features | Reference/source |
|---|---|---|
| pUC18 | Cloning vector, $Amp^R$ | Thermo Fisher Scientific |
| pCR2.1-TOPO® | TA cloning vector, $Amp^R$, $Kan^R$ | Thermo Fisher Scientific |
| pCR2.1/ΔCj1608 | pUC18 derivative containing *Cj1608* flanking regions and *aphA-3* for allelic exchange of *Cj1608* for *aphA-3* | This study |
| pTZ57R/TΔHP1021 | pTZ57R/T derivative containing *aphA-3* gene | [4] |
| pUC18/COM/Cj1608 | pUC18 derivative containing *Cj1608* flanking regions and *Cj1608* for allelic exchange of *aphA-3* for *Cj1608* | This study |
| pSB3021 | Suicide vector for integration of *cat* gene in *hsdM* gene for complementation | [5] |
| pUC18/ΔAbu0127 | pUC18 derivative containing *Abu0127* flanking regions and *aphA-3* for allelic exchange of *Abu0127* for *aphA-3* | [6] |
| pET28/StrepCj1608 | pET28Strep derivative containing the *Cj1608* gene for protein expression | This study |
| pET28/StrepAbu0127 | pET28Strep derivative containing the *Abu0127* gene for protein expression | This study |

| Recombinant protein | Relevant features | Reference/source |
|---|---|---|
| StrepCj1608 | Recombinant, *C. jejuni* Cj1608 protein, Strep-tagged at N-terminus, purified from *E. coli* | This study |
| StrepAbu0127 | Recombinant, *A. butzleri* Abu0127 protein, Strep-tagged at N-terminus, purified from *E. coli* | This study |

**Table S2: Primers used in this study.**

| Oligo name | Sequence (5' → 3') |
|---|---|
| **P1** | gatcaaaacctagaatttacagatg |
| **P2** | cacccgggtaccgagtcataatcttgtccaatcaaaatatt |
| **P3** | tggacaagattatgactcggtacccgggtgactaa |
| **P4** | ctgatttcatatttcttcctttctaaaacaattcatccagtaaaatatag |
| **P5** | ctatattttactggatgaattgttttagaaaggaagaaatatgaaatcag |
| **P6** | ctgctttatcaatagtccaacg |
| **P7** | cagatacaacactttttgacaatg |
| **P8** | gctcgacatactgttcttccc |
| **P9** | acagctatatccagcatcacttag |
| **P10** | gtcgactctagaggatccccggatttaaagcacatagtgatgg |
| **P11** | ctcctagttagtcaggatcctcatcttttttccaattataatat |
| **P12** | ggatcctgactaactaggaggaataaatg |
| **P13** | ctaaaacaattcatccagtaaaat |
| **P14** | ttttactggatgaattgttttaggggataatgtatgaatttgagaaa |
| **P15** | cgaattcgagctcggtacccccctgcaataatatcatcaccc |
| **P16** | ttttgacattcatgcctttgaagag |
| **P17** | gataaaccgcagctgttgcaatt |
| **P18** | gtttgttttcagcaagccactc |
| **P19** | gccaagcgttgtagatatgtca |
| **P23** | aaaccaccagaacaggcaca |
| **P24** | tgacgtttcaagatttggagcag |
| **P25** | taccataacccatagcaccgat |
| **P26** | agaaattgaaggcgatatgggc |
| **P27** | tctggaccttgaacaaattgca |
| **P28** | agccttcggttcttcctaca |
| **P29** | cgggatccatgaaagttttaattattgaaaatg |
| **P30** | ccggtcgacttatttttttcttgctgatttcata |
| **P31** | cgggatccatgaacatattaattatcgaaaatg |
| **P32** | ccggtcgacctatttcttttttcgatatctag |
| **P33** | ggagtaagaatagcttcgaatggcccagttgtaccgtcatata |
| **P34** | ctttattcagcgcttattttaacagc |
| **P35** | ggagtaagaatagcttcgaatgaatagctgctcttgaagcagc |
| **P36** | cgctatcaaataccatgcgac |
| **P37** | ggagtaagaatagcttcgaatcaatgaaaattgtggaagct |
| **P38** | agattgaatacagctttgagt |
| **P39** | ggagtaagaatagcttcgaattccacttgttataagttcatcattt |
| **P40** | acttgatattagaagaattgcaact |
| **P41** | FAM-ggagtaagaatagcttcgaat* |
| **P42** | Cy5- ggagtaagaatagcttcgaat** |
| **P43** | cggaatttgacattttcgctcc |
| **P44** | gcggttatttgaggctattattgttatga |
| **P45** | gtggaagctaatattaattccgttgaca |
| **P46** | tgtttcttttggtagcaacgga |

* - FAM, 6-fluorescein amidite;

** - Cy5, tetramethylindo(di)-carbocyanines 5.
